# Draft genome sequences of *Hirudo medicinalis* and salivary transcriptome of three closely related medicinal leeches

**DOI:** 10.1101/357681

**Authors:** Vladislav V. Babenko, Oleg V. Podgorny, Valentin A. Manuvera, Artem S. Kasianov, Alexander I. Manolov, Ekaterina N. Grafskaia, Dmitriy A. Shirokov, Alexey S. Kurdyumov, Dmitriy V. Vinogradov, Anastasia S. Nikitina, Sergey I. Kovalchuk, Nickolay A. Anikanov, Ivan O. Butenko, Olga V. Pobeguts, Daria S. Matushkina, Daria V. Rakitina, Elena S. Kostryukova, Victor G. Zgoda, Isolda P. Baskova, Vladimir M. Trukhan, Mikhail S. Gelfand, Vadim M. Govorun, Helgi B. Schiöth, Vassili N. Lazarev

## Abstract

Salivary cell secretion (SCS) plays a critical role in blood feeding by medicinal leeches, making them of use for certain medical purposes even today. We annotated the *Hirudo medicinalis* genome and performed RNA-seq on salivary cells isolated from three closely related leech species, *H. medicinalis, Hirudo orientalis,* and *Hirudo verbana.* Differential expression analysis verified by proteomics identified salivary cell-specific genes, many of which encode previously unknown salivary components. However, the genes encoding known anticoagulants were not differentially expressed in the salivary cells. The function-related analysis of the unique salivary cell genes enabled an update of the concept of interactions between salivary proteins and components of haemostasis. Thus, our study provides one of the most comprehensive knowledge of the genetic fundamentals of the blood-sucking lifestyle in leeches.

## Introduction

The genome sequencing of haematophagous animals and transcriptional profiling of their salivary glands has attracted considerable attention in recent years because many haematophagous species transmit various infectious diseases caused by viruses, bacteria, protozoa, and helminths. The elucidation of the genetic mechanisms that allow haematophagous species to act as vectors of pathogenic organisms [1–5] is of great importance for public health care, veterinary medicine, and microbiology. Opposite to other hematophagous species, blood-sucking leeches, belonging to the subclass *Hirudinea* (true leeches) of the phylum *Anelidae,* attract interest in regard to the identification of novel bioactive compounds. Blood-sucking leeches and, in particular, medicinal leeches, have been used for bloodletting to treat diverse ailments since ancient times [6]. Although the use of live leeches to treat human diseases is not encouraged by current medicine because of the high risk of an undesired outcome, hirudotherapy is still indicated in certain medical conditions. In particular, application of medicinal leeches improves tissue drainage after replantation when the common surgical correction of venous congestion fails or is unfeasible [7]. In these cases, hirudotherapy frequently provides beneficial effects because the leech feeding apparatus has evolved to promote the finely tuned inhibition of haemostasis and blood coagulation [8, 9]. The composition of leech saliva has been shown to play a key role in this inhibition [9, 10].

In medicinal leeches, which belong to the group of so-called *jawed leeches,* saliva is secreted by unicellular salivary glands that reside in the anterior part of the body and are interspersed between the muscle fibres that connect the jaws with the body wall. Each salivary cell extends a single duct from the cell body to the jaw, and the duct ends in a tiny opening between the calcified teeth of the jaw. Medicinal leeches incise the host skin at the feeding site by their jaws and release the salivary cell secretion (SCS) into the wound [9]. During one act of blood meal, medicinal leeches partially or completely empty their salivary gland cells [11]. However, salivary gland cells replenish their reservoirs with the content of SCS within seven days after blood meal, making a leech to be got ready for another act of blood feeding [12]. In addition to its known inhibitory effect on haemostasis and blood coagulation, SCS suppresses inflammation, exhibits analgesic effects, possesses antimicrobial activity, and alters vasodilatory responses, enhancing local blood circulation to facilitate leech feeding. Moreover, some SCS components are thought to preserve the blood from rapid degradation after ingestion.

Although transcriptional analysis of the salivary glands of jawless and jawed leeches was attempted [13–16] and some SCS components have been characterized and their respective targets in hosts have been identified [9], the repertoire of the bioactive saliva components remains largely unknown. Elucidating the SCS composition in medicinal leeches is the key to understanding (i) the molecular mechanisms underlying the orchestrated interaction between the leech SCS and the components of haemostasis, (ii) evolution of the blood feeding lifestyle in leeches, and (iii) genetics of hematophagy. Identification of the unknown bioactive SCS components will facilitate the development of novel pharmacological compounds for treating impaired peripheral blood circulation, venous congestion or microbial infections.

In the current study, we performed sequencing, genome assembly and annotation *H. medicinalis* genome as well as transcriptional profiling of the salivary cells followed by proteomic validation of SCSs of three medicinal leeches, *Hirudo medicinalis, Hirudo orientalis,* and *Hirudo verbana.* This study aims to provide a comprehensive map of the genetically encoded components of blood meal-related genes in leeches.

## Results

### Genome assembly and annotation

To assemble the *H. medicinalis* genome, we extracted DNA from an adult leech. Before being processed, the leach was maintained without feeding for at least two months. We created a set of three shotgun libraries to perform sequencing by using three different platforms (**Supplementary Table 1**). All read datasets were combined, and a single assembly was created by SPAdes [17]. The resulting assembly contained 168,624 contigs with an N50 contig length of 12.9 kb (**Supplementary Table 2**).

Preliminary analysis (contigs Blast) revealed the presence of bacterial sequences in the resulting assembly. Therefore, we conducted binning to discriminate the leach contigs (a leech bin). We built a distribution of contigs according to their GC abundance, tetranucleotide frequencies, and read coverage. To increase the binning accuracy, the read coverage was determined by combining the DNA reads with the reads corresponding to a combined transcriptome of *H. medicinalis* (see below). The discrimination of the eukaryotic and prokaryotic contigs is illustrated in **Fig. 1A/B, Supplementary Table 3** and **Supplementary Data 2**. Additionally, we selected the mitochondrial contigs to assemble the leech mitochondrial genome [18].

The eukaryotic contigs underwent a scaffolding procedure using paired reads. Scaffolds were generated using Illumina paired-end and mate-pair read datasets by SSPACE [19]. After scaffolding, the assembly consisted of 14,042 sequences with an N50 scaffold length of 98 kb (**Supplementary Tables 4 and 5**). The total length of the genome draft was estimated to be 187.5 Mbp, which corresponds to 85% of the theoretical size of the leech genome (**Supplementary Table 6**). A total of 14,596 protein coding genes were predicted.

**Fig. 1.**
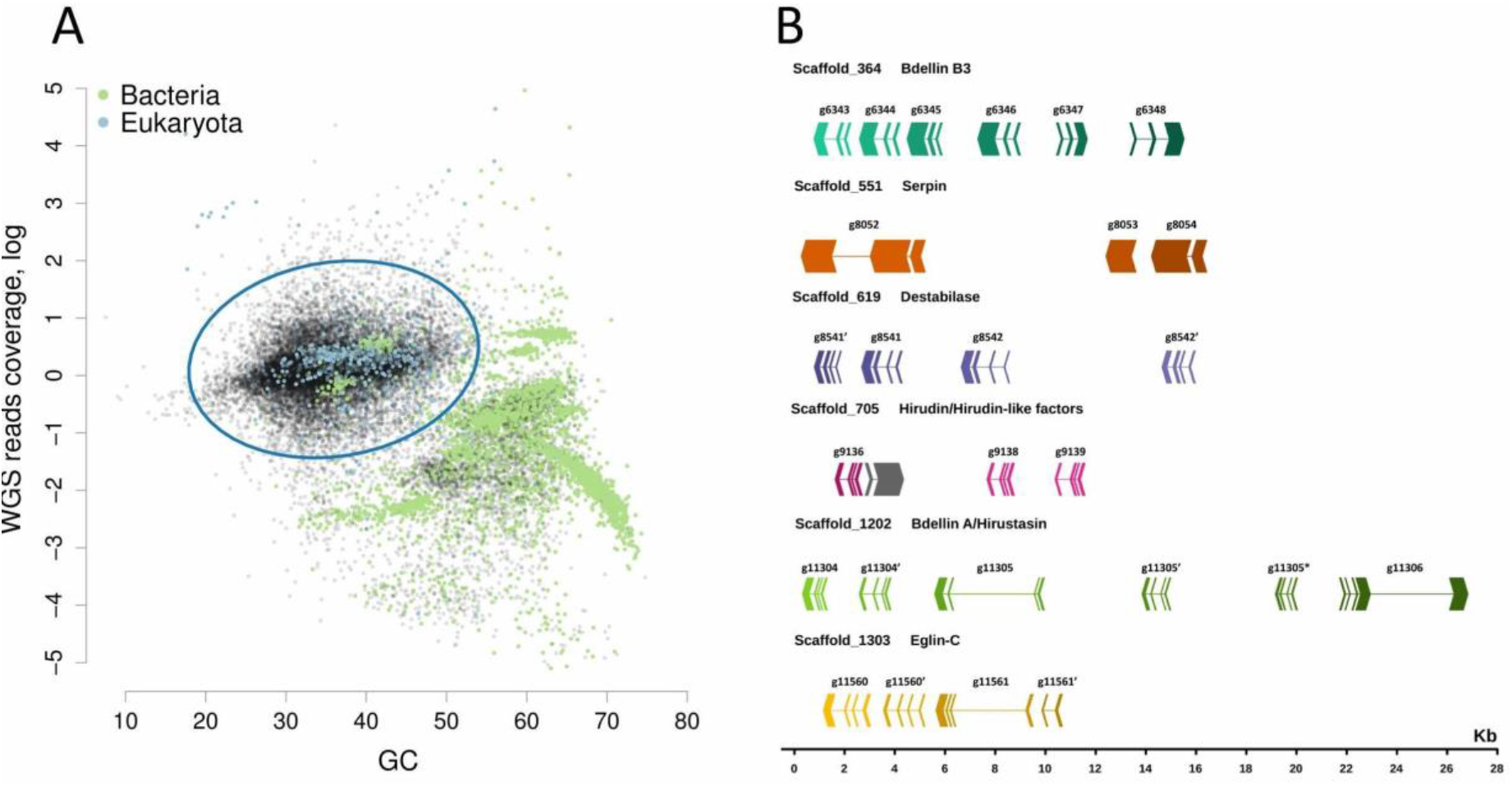
The *H. medicinalis* genome binning. **(A)**. 2D-plot showing the contig distribution in coordinates of GC content and coverage by a combination of reads obtained by Ion Proton and Illumina. Contigs are indicated by dots, and the taxonomic affiliation of contigs at the domain level is encoded by colour (green – *Bacteria,* blue – *Eukarya,* black – no assignment). The taxonomic affiliation was determined by direct Blastn search against the National Center for Biotechnology Information (NCBI) nt database. The 3D plot showing the contig distribution in coordinates of GC content, read coverage (Proton and Illumina), and host cDNA read coverage is presented in **Supplementary Data 1**. **(B)** *H. medicinalis* genome contains clusters of blood meal-related genes. The graph shows the exon-intron structure of genes and arrangement of gene clusters in scaffolds on a general scale. The exon arrows indicate the direction of transcription (gray – unknown gene).

Also, we identified new homologs of genes encoding known anticoagulants or blood meal-related proteins. The multiple amino acid alignments for each of these protein families (**Supplementary Figs. 1,2**) Based on the genome sequence data and using known protein sequences, we determined the organization of these genes (**Supplementary Table 7, Fig1 B**.). Positions and lengths of exons and introns were predicted using the respective cDNA and protein sequences as references. In some cases, genes are localized in common scaffolds and form tandems or clusters **Fig1 B**.

### mRNA-seq, transcriptome assembly and annotation

To obtain tissue-specific mRNA samples from three medicinal leech species, *H. medicinalis, H. verbana,* and *H. orientalis,* we isolated salivary cells and muscles from the cryosections of the anterior body parts using laser microdissection (**Fig. 2A**). Then, we constructed two cDNA libraries with and without normalization for each mRNA sample using the oligo-dT primer and sequenced them on the Ion Torrent PGM (**Supplementary Table 8**). Four read datasets corresponding to the constructed cDNA libraries were used for the *de novo* assembly of a combined transcriptome for each medicinal leech species using the Trinity RNA assembler [20] (**Supplementary Table 9**). We used the combined transcriptomes to map non-normalized tissue-specific reads. Read mapping was necessary to perform consecutive differential expression analysis.

Gene Ontology (GO) analysis of the detected transcripts was performed using Blast2GO [21] and BlastX. The ‘nr’ database served as a reference database. GO analysis demonstrated that all three medicinal leech species had similar transcript distributions across GO categories (**Supplementary Figure 3**). The taxonomy distribution of the closest Blast hits also was similar (**Supplementary Figure 4**). The majority of the identified transcripts were found to match two species of *Annelida:* 59.8% to *H. robusta* and 10.7% to *C. teleta.* This analysis also confirmed the absence of contamination by non-leech transcripts.

The prediction of coding regions (or open read frames, ORFs) and annotation of transcriptomic data were carried out using Transdecoder and Trinotate. ORFs were translated using the BlastP algorithm, and the protein sequences were annotated by EuKaryotic Orthologous Groups (KOG) classification using the eggNOG database [22] (**Supplementary Figure 5**). The KOG classification revealed that all three medicinal leech species have similar transcript distributions across KOG categories. All three medicinal leech species were also found to share the vast majority of their orthologous clusters (**Supplementary Figure 6**).

**Fig. 2.**
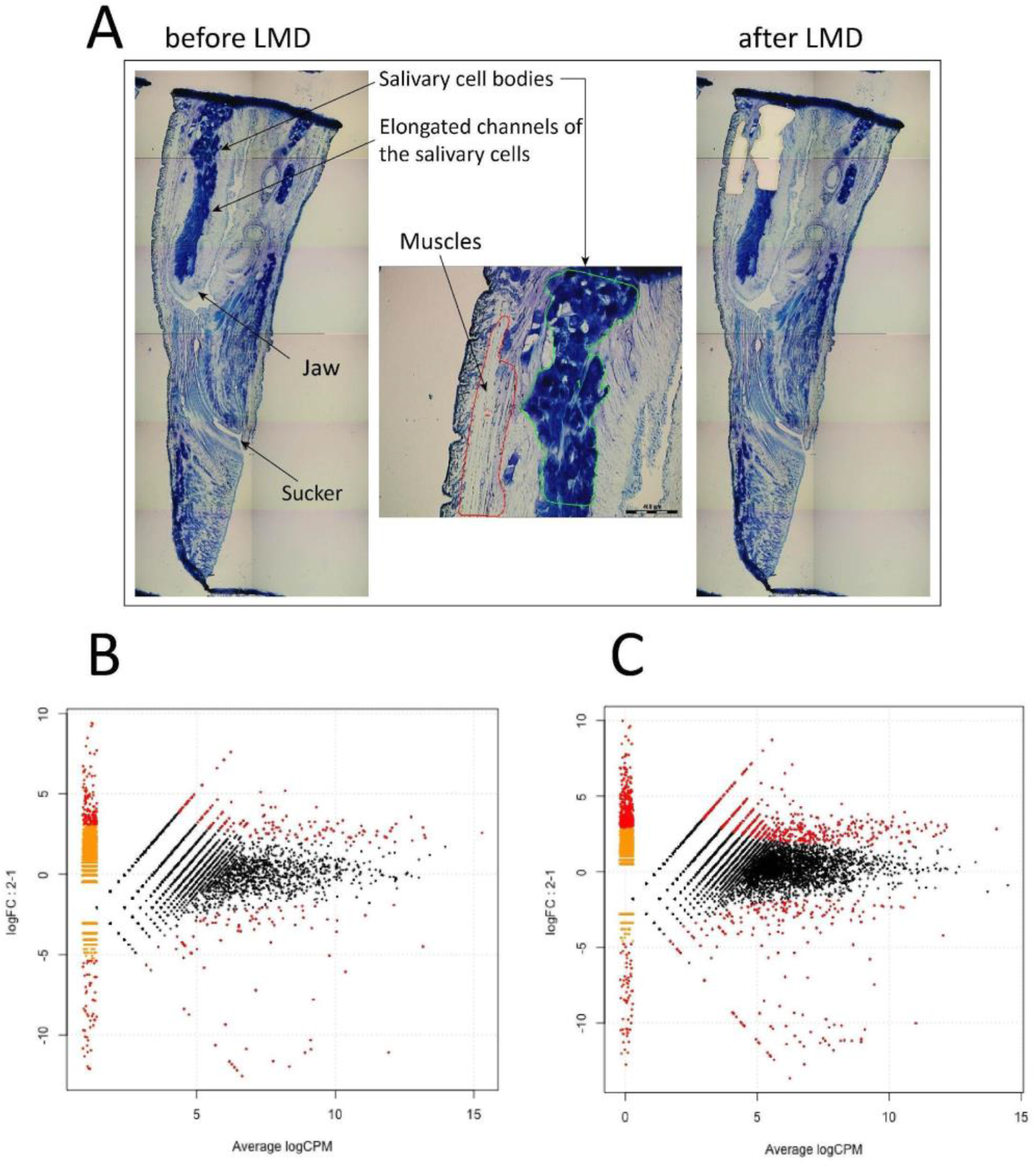
Differential expression analysis of salivary cells. **(A)** Isolation of salivary cells and muscles by laser microdissection. MA plots of differentially expressed genes in the salivary cells and muscles of *H. medicinalis* for the *de novo* assembled transcriptome **(B)** and the genome model **(C)**.

### Differential expression analysis

To estimate the relative expression levels of the transcripts identified in the salivary cells and muscles and to identify transcripts unique to the salivary cells, we mapped the tissue-specific cDNA reads without normalization against the combined transcriptome of each medicinal leech species. We also mapped the tissue-specific cDNA reads of *H. medicinalis* against its genome assembly. Differentially expressed genes were detected according to a recent protocol [23]. To identify genes that are differentially expressed in the salivary cells and muscles, an individual MA plot was constructed for each medicinal leech species using its combined transcriptome (**Fig. 2B, Supplementary Figure 7**). An additional MA was constructed for *H. medicinalis* using its genome assembly (**Fig. 2C**). Genes with a q-value (FDR) < 0.05 were considered to be differentially expressed.

We identified 102, 174, and 72 differentially expressed transcripts in the salivary cells of *H. medicinalis, H. orientalis,* and *H. verbana,* respectively. Because the three are closely related medicinal leech species, the protein sequences of the differentially expressed transcripts were grouped into orthologous clusters to simplify the subsequent functional analysis. We identified 25 differentially expressed, orthologous clusters shared by three leech species and 44 orthologous clusters shared by at least two leech species (**Fig. 3, Supplementary Tables 10-11**). The majority of sequences in the identified orthologous clusters correspond to hypothetical proteins annotated in the genome of *H. robusta*. Analysis of conserved domains in the identified orthologous clusters allowed the determination of sequences belonging to known protein families.

We also analysed the differentially expressed genes of *H. medicinalis* using its genome assembly. The cDNA reads for the salivary cells, muscles, and neural tissue [24] (reads were obtained from the Sequence Read Archive (SRA)) were mapped onto the genome assembly. For the neural tissue, we used a read dataset for ganglion 2 because of its localization in the preoral segments. Differential expression analysis identified 42 genes unique to the salivary cells of *H. medicinalis* (**Supplementary Table 12**).

### Proteomics of salivary cell secretion

For proteomic analysis, we collected SCSs from three medicinal leech species, *H. medicinalis, H. orientalis,* and *H. verbana,* which were maintained without feeding for at least two months. The SCSs were collected according to a previously reported method [25] with some modifications (see Methods).

The sample preparation method is critical to the resultant repertoire of the identified proteins because the SCS consists of both low- and high-molecular-weight components [9] and contains proteinase inhibitors, glycoprotein complexes, and lipids. The latter may form complexes with proteins [26]. Therefore, we combined several sample preparation methods and several mass spectrometry techniques to cover the broadest repertoire of the SCS proteins. Proteomic datasets obtained by different sample preparation methods and mass spectrometry techniques were combined to create a final list of the identified proteins for each medicinal leech species.

We identified 189, 86, 344 proteins in the SCSs of *H. medicinalis, H. orientalis,* and *H. verbana,* respectively and grouped them into orthologous clusters as described above. All three medicinal leech species were found to share 39 orthologous clusters, and 50 orthologous clusters were shared by at least two species (**Fig. 3, Supplementary Table 13**). Combination of the transcriptomic and proteomic data revealed 25 orthologous clusters of genes unique to the salivary cells (**Supplementary Table 11**). A list of individual components of the leech SCS is given in **Fig. 3**. Surprisingly, the expression of genes encoding known SCS anticoagulants and blood meal-related proteins did not differ in salivary cells and in muscles. To validate this finding, we examined the expression of saratin, eglin C, bdellins, hirustasin, destabilase, metallocarboxypeptidase inhibitor, apyrase, and angiotensin converting enzyme (ACE) by the real time PCR of additional, independent tissue-specific cDNA libraries constructed for salivary cells and muscles. The real-time PCR results (data not shown) confirmed this finding. This indicates that genes encoding anticoagulants and blood meal-related proteins are involved not only in the blood feeding, but contribute to other, yet unknown physiological functions.

**Fig. 3.**
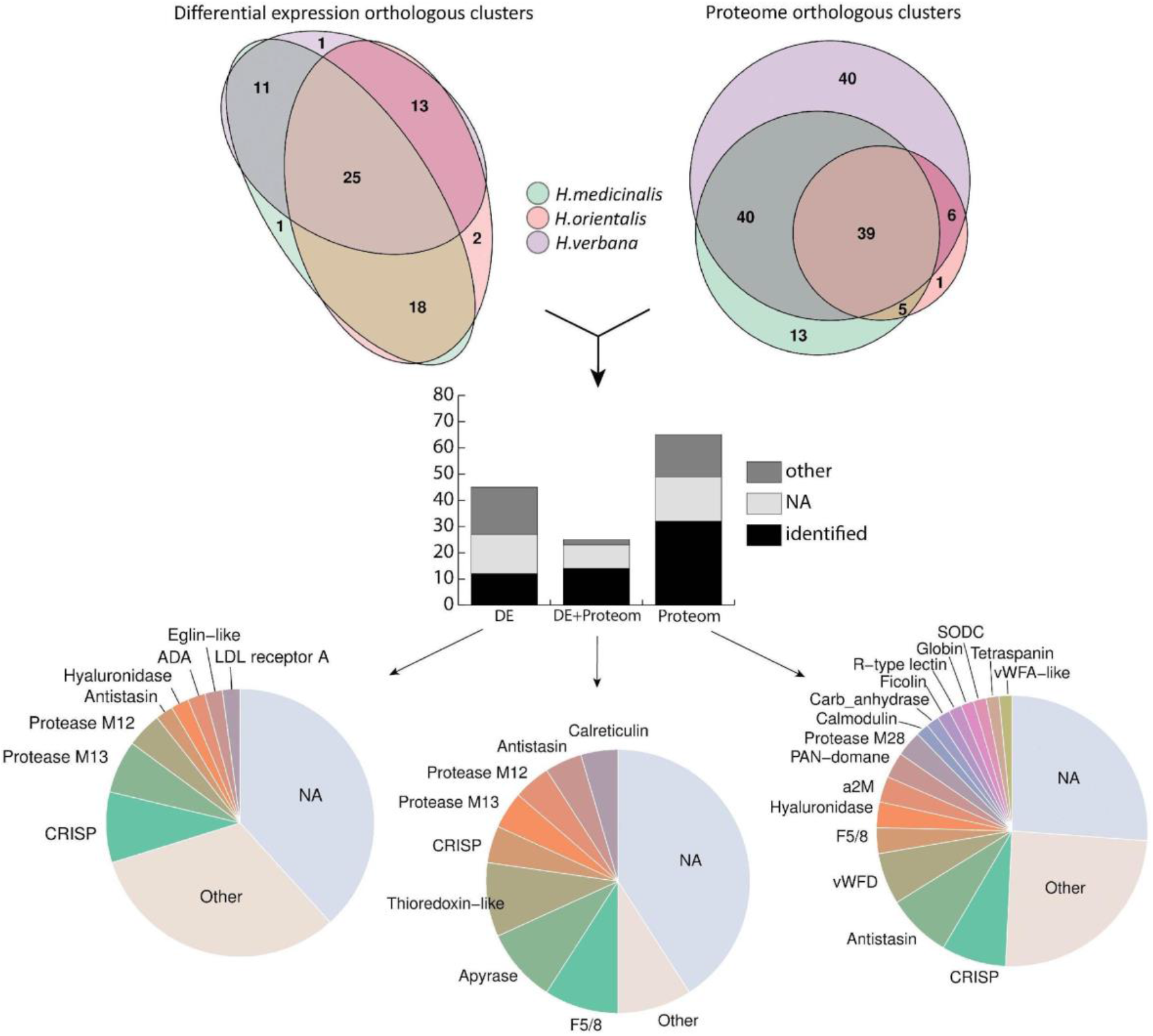
Summary of the identified SCS components. The Venn diagrams in the upper panel show the numbers of ortholog clusters identified by differential expression (DE) and proteomic (Prot) analyses across three medicinal leech species. The histogram in the middle panel features the numbers of orthologous clusters identified by the differential expression analysis, proteomic analysis or a combination thereof (DE+Prot). Each bar consists of ortholog clusters identified as known blood feeding-related components (identified), other known proteins (other), and unknown proteins (NA). The pie charts in the lower panel illustrate the abundance of the individual SCS components identified by the differential expression analysis, proteomic analysis or their combination. For details, see **Supplementary Tables 11, 12, and 14**.

Below, we characterize SCS components classified into functional groups and describe their possible roles in the hemostasis. The sequences of proteins and their alignment are presented in **Supplementary Figs. 8-23**.

### Enzymes

#### Proteases

The results of this study show that metalloproteases of the M12, M13, and M28 families are the major enzymatic components of the SCS. The M12B (ADAM/reprolysin) peptidases are a large family of disintegrin-like metalloproteinases that have a broad range of functions and are involved in many physiological processes [27]. These enzymes are often found in snake venoms while the transcripts are observed in sialotranscriptomes of various hematophagous species [28–30]. In haemostasis, secreted proteases of the M12 family can participate in the inhibition of platelet adhesion [31, 32] and in clot softening due to the degradation of fibrinogen. These proteins exhibit metal-dependent proteolytic activity against extracellular matrix proteins (gelatine, fibrinogen, fibronectin), thereby affecting the regulation of inflammation and immune responses.

In mammals, proteases of the M13 family are involved in the formation and development of the cardiovascular system and in the regulation of neuropeptides in the central nervous system [33]. One of their most important functions is the activation of biologically active peptides, particularly peptides involved in the regulation of blood pressure (angiotensin and bradykinin). In mammals, ACE is an important component of the renin angiotensin system (RAS). ACE is expressed in the sialotranscriptomes of the leech *(Theromyzon tessulatum),* the cone snail *(Conidae),* the vampire snail *(Colubraria reticulata),* and dipteran species *(Diptera)* [34, 35].

The identified sequences of M28 family exopeptidases belong to the Q-type carboxypeptidases, also known as lysosomal dipeptidases or plasma glutamate carboxypeptidase (PGCP). These peptidases were shown to be involved in the regulation of the metabolism of secreted peptides in the blood plasma and the central nervous system in mammals [36]. These enzymes appear to serve to deactivate certain signalling peptides in the blood and are components of haemoglobinolytic systems in haematophagous parasites, playing the role of digestive exopeptidases [37]. Notably, leech salivary gland secretions contain carboxypeptidase inhibitors, which presumably prevent the untimely digestion of the blood meal by other types of peptidases [9, 38].

#### Superoxide dismutase (EC 1.15.1.1)

We identified sequences of secreted superoxide dismutase family (SODC, Cu/Zn type) enzymes. This family of metalloproteins is mainly typical of eukaryotes and is involved in free radical inactivation, which retards oxidative processes. In the blood, superoxide dismutase catalyses the conversion of superoxide into molecular oxygen and hydrogen peroxide and prevents the formation of peroxynitrite and hydroxyl radicals [39]. Interestingly, peroxynitrite may suppress haemostatic function by the nitration of key procoagulants [39, 40], while hydrogen peroxide is a key signalling molecule involved in the regulation of many processes (coagulation, thrombosis, fibrinolysis, angiogenesis, and proliferation). In ticks, SODC is presumed to participate in regulating the colonization of the intestinal tract by bacteria, including causative agents of diseases [41]. In SCSs, SODC appears to exhibit an antibacterial effect along with other proteins of the innate immune system and prevents unwanted blood oxidation during feeding and digestion. Notably, haem-containing compounds and free iron are involved in the formation of free radicals and the provocation of oxidative stress [42].

#### Carbonic anhydrase (EC 4.2.1.1)

This enzyme is a key component of the bicarbonate buffer system and is involved in the regulation of pH values in the blood, the digestive tract, and other tissues [43, 44]. In haematophagous animals, this enzyme can maintain optimal conditions for the digestion of a blood meal [45, 46]. Carbonic anhydrase appears to cause a local increase in acidosis at the bite site, decreasing the activity of blood coagulation factors.

#### Hyaluronidase (EC 3.2.1.35)

These enzymes are common in the proteomic and transcriptomic data of haematophagous and venomous animals. The salivary secretions of different leech species are known to contain hyaluronidase (heparinase, orgelase) [47]. In the proteome and transcriptome, we found three clusters containing a domain of the glycosyl hydrolase family 79 (O-glycosyl hydrolases). This family includes heparinases, which play an important role in connective tissues. In venoms and salivary gland secretions, these enzymes catalyse the hydrolysis of hyaluronic acid, resulting in the loss of structural integrity of the extracellular matrix and thereby facilitating the penetration of anticoagulants and other active molecules deeper into the tissues [48]. In addition, the low-molecular-weight heparin produced by cleavage suppresses and inhibits blood coagulation [49].

#### Apyrase (EC 3.6.1.5)

Apyrases are nucleotidases involved in the enzymatic degradation of ATP and ADP to AMP. Secreted apyrases and 5’-nucleases are well-known and well-characterized components of the salivary gland secretions of venomous and haematophagous animals, including leeches [10]. Apyrases are anticoagulants because they remove ADP, an important inducer of platelet aggregation at sites of tissue injury [50].

#### Adenosine/AMP deaminase (EC:3.5.4.4)

Adenosine deaminase (ADA) catalyses the hydrolytic deamination of adenosine to form inosine. Adenosine deaminases are well studied and have been found in the saliva of various blood-sucking insects [51]. ADA is also found in the salivary gland secretion of the vampire snail *C. reticulata,* which belongs to *Spiralia,* as well as leeches [52]. ADA is thought to play an important role in the removal of adenosine because of its involvement in pain perception processes [53].

### Proteinase inhibitors

#### Antistasins

We identified sequences corresponding to proteinase inhibitor I15 (leech antistasin) **Fig.4.** Proteins of this family are commonly found in blood-sucking leeches and play a key role in the inhibition of blood coagulation. Their main targets are serine proteases participating in haemostasis, such as the factor Xa, kallikrein, plasmin, and thrombin [9]. Decorsin, an antistasin from *Macrobdella decora,* was demonstrated to inhibit platelet aggregation [54], and gigastasin from the giant Amazon leech *(Hementaria ghilianii)* was recently reported to potently inhibit complement C1 [55]. Antistasin from *Hementeria officinalis* is the closest homologue of the sequences identified in our study.

**Fig. 4.**
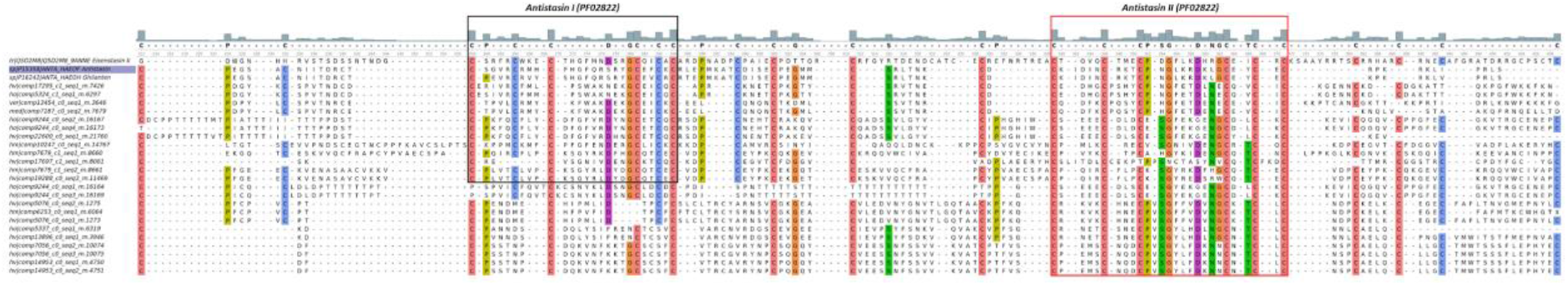
Multiple sequences alignment of Antistasin-like transcripts with dual domain antistasin-type protease inhibitors from leeches’ Antistasin *(Haementeria officinalis,* P15358), Ghilantein *(Haementeria ghilianii,* P16242) and Eisenstasin II from earthworm *(Eisenia andrei,* Q5D2M8J. The boxes indicates two antistasins domane. Alignment is generated by MUSCLE algorithm, residues are colored according to ClustalX colour scheme, conserved amino acids are colorized by conservation level (threshold > 50%). Reference sequence are marked purple.

#### CAP/CRISP

The cysteine-rich secretory protein/antigen 5/pathogenesis-related 1 proteins (CAP) superfamily includes numerous protein families, particularly cysteine-rich secretory protein (CRISP) **Fig 5A**. They are commonly found in the venoms of snakes and other reptiles, and most of them are toxinsy [56, 57]. In some investigations, CRISPs from haematophagous species were thought to be involved in haemostasis (HP1). The identified sequences show similarity to protein sequences from the haematophagous parasitic nematode *Ancylostoma caninum* (hookworm), such as the potassium channel blocker AcK1 [58] and the possible platelet aggregation inhibitor HPI [59], as well as to the snake toxins triflin *(Protobothrops flavoviridis)* and natrin-1 *(Naja atra)* [60, 61]. Among the differentially expressed genes, we identified sequences with a new “Cys-rich” motif **Fig 5B**. This group of proteins is characterized by the presence of a signal peptide and two cysteine patterns CX{5,14}CX{7}CX{8}CC{2}C and CX{7,17}CX{9}CX{8}CC{2}C.

**Fig. 5.**
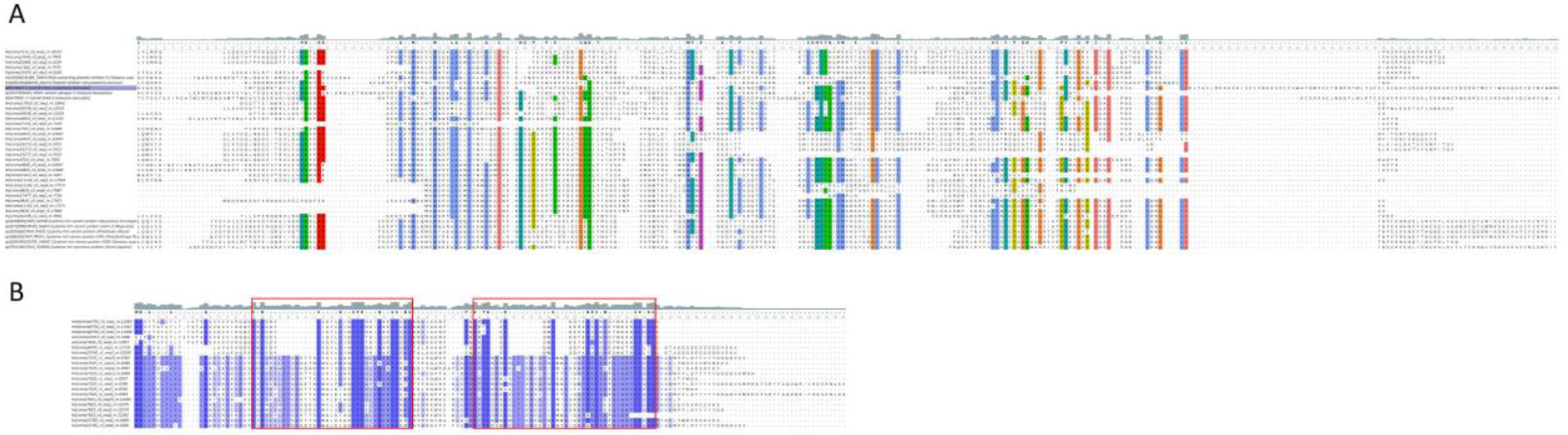
(A) Alignment of CRISP domains with diverse CAP/CRISP proteins. Putative platelet inhibitors from (Ancylostoma caninum, Q962V9), (Tabanus yao, C8YJ99), CAP domain containing proteins from Vampire Snail (Cumia reticulata, QBH70087.1; QBH70092.1) and reptile Cystein-rich venom proteins triflin (Protobothrops flavoviridis), natrin-2 (Naja atra) and other. Alignment is generated by MUSCLE algorithm, residues are colored according to ClustalX colour scheme, conserved amino acids are colorized by conservation level (threshold > 50%). Reference sequence are marked purple. (B) Alignment of new “Cys-rich” domains. The boxes indicates two cysteine patterns, amino acids are colorized by percentage Identity coloring scheme.

#### Eglin-like

Eglins are small cysteine-free proteins that belong to the I13 family of serine proteinase inhibitors [62]. Eglins from leeches have inhibitory activity against neutrophil elastases and cathepsins G and also to participate in the protection of the crop contents from untimely proteolysis [9]. Of note, sequences identified in the present study have low homology to the classical eglin from leech **Fig.6A**.

#### Cystatin

We identified a cystatin sequence only in the proteome of *H. verbana.* Cystatins are small protein inhibitors of cysteine proteases (cathepsins B, H, C, L, S) [63] and are often found in the sialotranscriptomes of various ticks [64]. In ticks, cystatins play an important role in processes related to immune response, the regulation of endogenous cysteine proteases involved in the digestion of blood and haem detoxification [65]. The nematode *Nippostrongylus brasiliensis* utilizes cystatins to evade the host immune system [66].

#### PAN domain

This domain is present in numerous proteins, including the blood proteins plasminogen and coagulation factor XI [67]. The PAN/apple domain of plasma prekallikrein is known to mediate its binding to high-molecular-weight kininogen, and the PAN/apple domain of the factor XI binds to the factors XIIa and IX, platelets, kininogen, and heparin [68]. The salivary gland secretion of the leech *H. officinalis* was found to contain the leech antiplatelet protein (LAPP), which has a PAN domain and is involved in haemostasis. This protein exhibits affinity for collagens I, III, and IV and thereby inhibits collagen-mediated platelet adhesion [69].

**Fig. 6.**
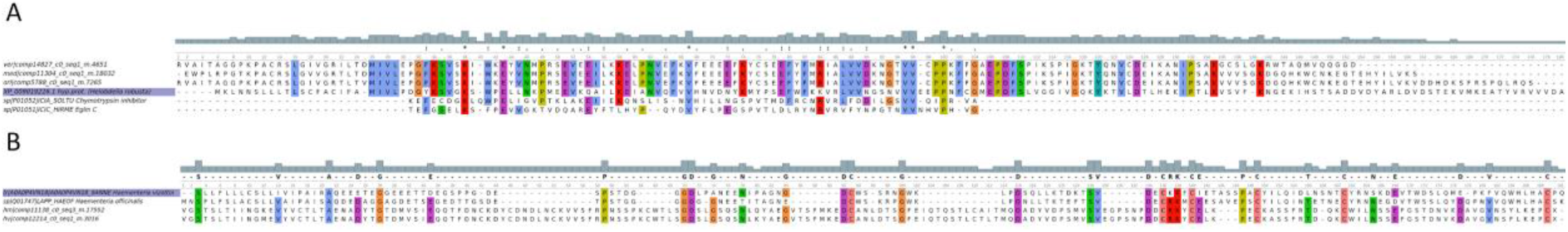
(A) Amino acid sequences alignment of Eglin-like transcripts with Eglin *(Hirudo medicinalis,* P01051), hypothetical protein *(Helobdella robusta,* xp_009019226.1) and chymotrypsin inhibitor homolog from Potato *(Solanum tuberosum,* P01052). Alignment is generated by MUSCLE algorithm, residues are colored according to ClustalX colour scheme. Identical and conserved residues indicated respectively by asterisk, period and colon. (B) Alignment of PAN domains with leech anti-platelet protein *(Haementeria officinalis,* Q01747) and putative anti-platelet-like protein *(Haementeria vizottoi,* A0A0P4VN18). Conserved amino acids are colorized by conservation level (threshold > 75%). Reference sequence are marked purple.

#### Alpha-2-macroglobulin (α2M)

The highly conserved, multifunctional α2M is involved in the inhibition of a broad range of proteases (serine, cysteine, aspartic, and metalloproteases), interacts with cytokines and hormones, and plays a role in zinc and copper chelation [70]. It can act as a plasmin inhibitor, thereby inhibiting fibrinolysis, but in some cases, it inhibits coagulation by inactivating thrombin and kallikrein [71]. This protein is believed not only to be involved in leech immune processes but also to be an important component of the salivary gland secretion that enhances anticoagulation processes.

### Molecules involved in adhesion

#### Ficolin

Ficolins are a component of the innate immune system and trigger a lectin-dependent pathway of complement activation [72]. In invertebrates, ficolins are involved in the recognition of bacterial cell wall components [73]. The fibrinogen-like domain is present in proteins with affinity for erythrocytes, *e.g.,* tachylectin-5A (TL5A). TL5A exhibits strong haemagglutinating and antibacterial activity in the presence of Ca^2+^ ions [74]. In reptile venoms, ficolin-like proteins, ryncolin [75] (from *Cerberus rynchops)* and veficolin-1 (UniProt: E2IYB3) (from *Varanus komodoensis),* are presumed to trigger platelet aggregation and blood coagulation.

#### F5/8 type C domain

A number of identified sequences contain one or several discoidin motifs (DS), known as the F5/8 type C domain. This domain is present in numerous transmembrane and extracellular proteins, *e.g.,* neuropilins, neurexin IV, and discoidin domain receptor proteins, and in proteins involved in haemostasis, such as the coagulation factors V and VIII [76]. The DS domain plays an important role in the binding of various ligand molecules, including phospholipids and carbohydrates [77]. Due to these features, DS-containing proteins are actively involved in cell adhesion, migration, and the proliferation and activation of signalling cascades [78]. Leech DS domain-containing proteins appear to act as lectins with high affinity to galactose and may be components of the innate immune system of the leech. In addition, they can bind to collagen or phosphatidylserine on the surface of platelets and the endothelium [79] and thus, by competitive inhibition, impair interactions between haemostatic factors.

#### Low-density lipoprotein receptor A family

The low-density lipoprotein receptor (LDLR) family is an important component of the blood plasma and is involved in the recognition and endocytosis of low-density lipoproteins in mammalian blood [80]. In contrast to known homologous proteins, these receptors are secretory rather than membrane proteins, and they contain four LDLR class A (cysteine-rich) repeats. Some invertebrates, including segmented worms, are hypothesized to be incapable of the synthesis of cholesterol and steroid hormones, and during feeding, leeches acquire cholesterol mainly from the blood of the host as an exogenous source [81]. We posit that this protein may be utilized by the leech for the scavenging and transportation of cholesterol-rich lipoprotein complexes.

#### R-type lectin

Proteins that contain the ricin-type beta-trefoil lectin domain have been found in prokaryotes and eukaryotes. In animals, R-type lectins exhibit diverse activities [82]. They are present in scavenger receptors (mannose, fucose, collagen receptors), N-acetylgalactosaminyltransferases, haemolytic toxins (CEL-III from *Cucumaria echinata)* and apoptosis-inducing cytotoxins [82, 83]. Previously, similar sequences were identified in leech transcriptomes; however, the authors assumed that this molecule has a mitochondrial localization [84]. Yet another noteworthy close homologue is the galactose-binding lectin EW29 from the earthworm *Lumbricus terrestris.* EW29 consists of two homologous domains and has been experimentally demonstrated to exhibit haemagglutinating activity [85]. As many known R-type lectins are involved in adhesion and trigger haemolysis [82], this molecule is of interest for further study.

#### vWFA domain

This domain is present in various plasma proteins: complement factors, integrins, and collagens VI, VII, XII, and XIV [86]. One protein identified in the leech proteome is a secreted protein that consists of four copies of the *vWFA* domain **Fig. 7**. The sequence contains several putative recognition sites: the metal ion-dependent adhesion site (MIDAS), the integrin-collagen binding site, and the glycoprotein Ib (GpIb) binding site. According to blast analysis, this domain is homologous to type VI collagen. Considering the domain organization of the protein and the presence of glycoprotein and collagen binding sites, one of the putative mechanisms of action involves binding to the surface of the endothelium or platelets, thereby preventing their interaction with collagen. This binding underlies the competitive inhibition during haemostasis (platelet scavenging) [34].

**Fig. 7.**
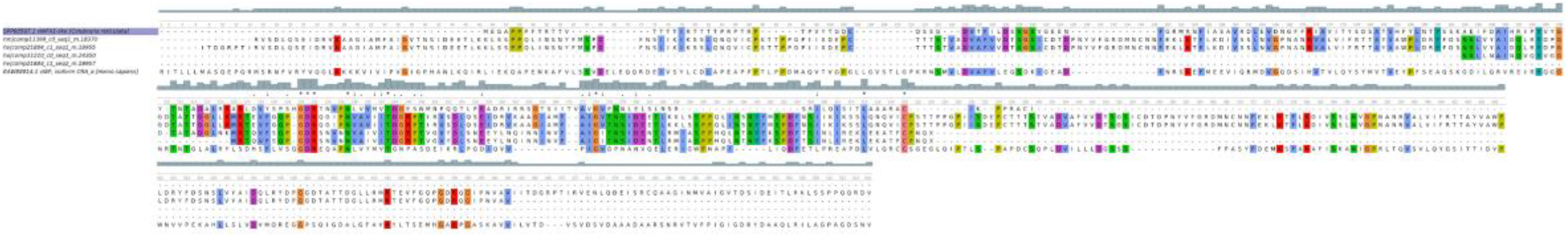
Alignment of the hirudo vWFA domains with the human vWFA1 (EAW88814.1) and vWFA1-like *(Colubraria reticulata,* SPP68597.1). Alignment is generated by MUSCLE algorithm, residues are colored according to ClustalX colour scheme. Identical and conserved residues indicated respectively by asterisk, period and colon. Reference sequence are marked purple.

## Discussion

Combined analysis of transcriptomic and proteomic data revealed that SCSs of three medicinal leech species, *H. medicinalis, Hirudo orientalis,* and *Hirudo verbena,* have a similar composition and to large extent contain many proteins homologous to those that are associated with either blood feeding in various hematophagous animals or have been identified in venoms of the poisonous organisms, such as M12 and M13 proteases, CRISP, Apyrase, ADA, cystatins, hyaluronidase and ficolins. However, we also detected novel salivary proteins, the roles and importance of which in blood feeding have yet to be established such as F5/8 type C domain, LDLR, PAN domain as well as proteins containing vWFD and vWFA domains. Moreover, we also found, somewhat surprisingly, that genes encoding well-characterised components of the medicinal leech SCS, such as hirudin, bdellins, eglins, saratins and destabilases were not differentially expressed in the salivary cells, suggesting that the proteins encoded by these genes support not only blood feeding but also some unknown functions in the leech.

We also identified new homologs of genes encoding known anticoagulants including antistasins and LAPP but also other blood meal-related proteins such as bdellins A and B3, eglin-C, hirustasin, destabilase, earatin/LAPP and leech DTI in the annotated genome. Interestingly, some of these genes are localized in common scaffolds and form tandems or clusters. This arrangement and the similarity of the genes suggest recent local duplications and many of them as likely to be linage specific. This unique diversity of anticoagulatant genes highlights their specific functional significance for the leech, which has led to their positive selection with subsequent expansion in the genome. It is also notable that almost all of these genes are serine protease inhibitors such as the antistasins, serpins, Kazal-Type Serine Protease Inhibitor, Potato inhibitor I family and hirudins. All of the above protein families are involved in host hemostasis or blood digestion [9, 84]. They are also the most broadly represented and diverse families in the salivary secret of leeches (**Supplementary Tables 10,11,13**).

It is possible that the initially duplication of some of these genes could be associated with dosage effect related to the amount of the gene product. During the divergence process, it is likely that some of these genes acquired increased specificity for new targets from the blood of various hosts. In the transcriptome data, we found that in most families of anticoagulants and products of blood meal related genes, the only one variant of the gene is expressed and these were the previously known sequences from the NCBI. At the same time, we also identified the expression of several variants of Bdellin B3, Lectin C-type, Destabilase, and three Elastase inhibitor genes (serpins). Gene duplication combined with positive selection is an important mechanism for the creation of new functional role for the existing genes (neofunctionalization) [87]. The functional segregation of hirudin and hirudin-like factor genes could be an example of this. The functions of hirudin are well understood, while targets of hirudin-like factors are still not known [88, 89]. Moreover, the results show that many well known blood meal-related genes are found in single-copies (without duplication) including Saratin, Leech DTI and Carboxy peptidase inhibitor. These gene families have probably very conserved functional activities in blood processing during feeding and we see no signs duplication of within these gene families.

For a long time, thrombin and factor Xa have been thought to be the main targets of components of the leech SCS. However, similar to those of other haematophagous species, the SCS of medicinal leeches were found to contain a complex mixture of molecules that inhibit both secondary and primary. Our findings lead to the suggestion that the action of the SCS components not merely on the key coagulation factors (thrombin and factor Xa) but on the different stages of haemostasis provides synergistic effects. Furthermore, functional analysis of the identified proteins demonstrated that leeches utilize ancient, highly conserved molecules as versatile affectors of the host haemostasis and immunity.

Similarity of the SCS compositions in the examined leeches with those of many other blood feeding species favours hypotheses about a convergent evolution of the saliva composition in evolutionary distant hematophagous animals as a result of having to adapt to the same targets in their hosts.

In the proteome of the SCSs, we found proteins that usually exhibit cytoplasmic or membrane localization, such as calreticulin, calmodulin, thioredoxin, chaperones, tetraspanin transcription factors, and certain ribosomal and cytoskeletal proteins. In contrast to that in jawless leeches [90], the mode of the secretion process in jawed leeches remains unknown. The presence of proteins with cytoplasmic or membrane localization appears however to be associated with apocrine secretion in the production of leech saliva.

Over all it is interesting that the SCS of the medicinal leeches have been found to contain proteins homologous to others found in a variety of the vertebrate species. These proteins are relatively conserved, and exhibit shared structural and functional features among various species. Some have been shown to be directly or indirectly involved in haemostasis in mammals. These proteins, avoiding interactions with components of the host immune system, could possibly affect the kinetics of biochemical reactions, thereby providing a synergistic effect of the SCS.

In sum, this analysis provides new insights into the role of the genome structure in the regulation of blood feeding-related gene expression and the evolutionary adaptation to the blood-sucking lifestyle. More broadly, the genome annotation performed in our study may serve as a blueprint for future experimentation on the medicinal leech as a model organism and provides a database of sequences encoding the unique bioactive leech proteins for use in developing novel pharmacological compounds.

## Methods

### BioProject and raw sequence data

The genome assembly was validated by the National Center for Biotechnology Information (NCBI). It was checked for adaptors, primers, gaps, and low-complexity regions. The genome assembly was approved, and the accession numbers MPNW00000000 and BioProject PRJNA257563 were assigned. All genome sequencing data were deposited in the Sequence Read Archive (SRA) with accession numbers (see **Supplementary Table 1**). Raw mRNA-seq data are available as FASTQ files and have been quality-checked and deposited in the SRA with their accession numbers (see **Supplementary Table 9**).

### Biological samples

Three leech species were provided by HIRUD I.N. Ltd. (Balakovo, Saratov Region, Russia). *H. medicinalis, H. verbana,* and *H. orientalis* were collected at a pond near Volkovo, Saratov region, Russia (51°91’03”, 47°34’90”), Lake Manych, Stavropol Krai, Russia (46°01’09”, 43°48’21”), and Lake Divichi, Kura-South Caspian Drainages, Azerbaijan (41°17’40”, 49°04’13”), respectively. Leech species were confirmed by sequencing the regions of the genes encoding nuclear and mitochondrial ribosomal RNA (for details, see **Supplementary Methods**).

### Genomic DNA extraction, WGS sequencing and genome assembly

Genomic DNA was extracted from a single adult leech *H. medicinalis* using the standard technique with slight modifications (for details see **Supplementary Methods**). The extracted DNA was purified (for details, see **Supplementary Methods**), and a set of three shotgun libraries was created.

#### Ion Proton shotgun sequencing

Pure genomic DNA (approx. 1000 ng) was fragmented to a mean size of 200-300 bp using the Covaris S220 System (Covaris, Woburn, Massachusetts, USA). Then, an Ion Xpress™ Plus Fragment Library Kit (Life Technologies) was employed to prepare a barcoded shotgun library. Emulsion PCR was performed using the One Touch system (Life Technologies). Beads were prepared using the One Touch 2 and Template Kit v2, and sequencing was performed using Ion Proton 200 Sequencing Kit v2 and the P1 Ion chip.

#### Ion Torrent mate-pair sequencing

A mate-pair library with 3-6 kb fragments was prepared from pure genomic DNA using Ion TrueMate Library Reagents (Life Technologies). The Ion PGM™ template OT2 400 kit (Life Technologies) was used to conduct emulsion PCR. Sequencing was performed by the Ion Torrent PGM (Life Technologies) genome analyser using the Ion 318 chip and Ion PGM™ Sequencing 400 Kit v2 (Life Technologies) according to the manufacturer’s instructions. To separate non-mate reads and split true mate reads, we used the matePairingSplitReads.py script.

#### Illumina mate-pair sequencing

A mate-pair library with 8-12 kb fragments was prepared from the pure genomic DNA using the Nextera Mate Pair Library Sample Prep Kit (Illumina) and TruSeq DNA Sample Prep Kit (Illumina) according to the manufacturer’s recommendations. The library was sequenced on the MiSeq platform (Illumina) using a 2 × 150 cycle MiSeq V2 Reagent Kit according to the standard Illumina sequencing protocols. Demultiplexing was performed using bcl2fastq v2.17.1.14 Conversion Software (Illumina). Adaptor sequences were removed from reads during demultiplexing. For trimming and separation of the single-end, paired-end, and mate-pair reads, NxTrim software was used.

Read datasets corresponding to the three shotgun libraries were combined, and SPAdes 3.6.0 software [17] was used to create a single genome assembly. For eukaryotic contig scaffolding, we used Sspace software [19] with the parameters -p 1 -x 0 -l library.txt -s Contigs.fasta -k 2.

### Contigs binning

#### Assembly analysis

The k-mer coverage and contig length were taken from the SPAdes assembly information (contig names). GC content was calculated using the infoseq tool built into the EMBOSS package. Contigs with a length less than 500 bp were ignored. The tetranucleotide content was calculated using <u>calc.kmerfreq.pl</u>, and the script is available at [https://github.com/MadsAlbertsen/miscperlscripts/blob/master/calc.kmerfreq.pl].

The reads of the individual shotgun libraries and the *H. medicinalis* cDNA reads (see below) were mapped against the genome assembly by bowtie2 [91]. The depth of read coverage for the individual libraries was calculated using BEDTools [92]. The taxonomic classification of the contigs was carried out by MEGAN6 [93] using the results of BLAST analysis (nr/nt database).

#### Classification of eukaryotic contigs

Using the R language and the car package, a concentration ellipse was built to confine at least 99% of those contigs against which at least ten cDNA reads had been mapped. To avoid the loss of potential eukaryotic contigs, we also considered the taxonomic affiliation of those contigs against which cDNA reads were not mapped. All contigs that were identified as neither prokaryotic nor eukaryotic but belonged to the ellipse, were assigned to be eukaryotic.

### Annotation of a draft genome

To annotate a draft genome, three sets of so-called hints, sequences in the genome that exhibit features of specific gene structures, such as exons, introns, etc., were generated (for details, see **Supplementary Methods**). The first set of hints was generated using sequences from the *H. robusta* protein coding genes. The second set of hints was generated using contigs corresponding to the *de novo* transcriptome assembly (see below). The third set of hints was generated using the cDNA reads (see below). All sets of hints were combined, and AUGUSTUS software [94] (version 3.7.1) was used for annotation of the draft genome.

### Laser microdissection

Laser microdissection was applied both to obtain tissue-specific mRNA samples from salivary cells and muscles for subsequent differential expression analysis and to collect different parts of the digestive tract (crop, coeca, intestine) for consecutive metagenomic analysis (for details, see **Supplementary Methods**). Briefly, live leeches were snap frozen in liquid nitrogen. The different parts of the leech bodies were cryosectioned, and slices were attached to membrane slides with the PEN foil (Leica Microsystems, Germany). Slides were stained with methylene blue to reveal salivary cell bodies. Staining was omitted when the parts of the digestive tract were isolated. Tissue collection was performed for three leech species, *H. verbana, H. orientalis* and *H. medicinalis,* by a Leica Laser Microdissection System LMD7000 (Leica Microsystems, Germany). Salivary cells and muscles were isolated directly into RNeasy extraction solution for the subsequent extraction of total RNA, and the parts of the digestive tract were isolated directly into ATL buffer from the QIAamp DNA Micro Kit for the subsequent extraction of total DNA.

### cDNA library construction and sequencing

Total RNA was extracted from the tissue fragments isolated by laser microdissection using an ExtractRNA Kit (Evrogen, Russia) (for details, see **Supplementary Methods**). Tissue-specific cDNA libraries were created using the Mint-2 cDNA Synthesis Kit (Evrogen, Russia) according to the manufacturer’s instructions. The adapters PlugOligo-3M and CDS-4M were used for cDNA synthesis. The normalization of cDNA was performed using the DNS (duplex-specific nuclease) in accordance with the manufacturer’s protocol (Evrogen, Russia). Normalized and non-normalized cDNA libraries (100 ng of each sample) were fragmented to a mean size of 400-500 bp by using a Covaris S220 System (Covaris, Woburn, Massachusetts, USA). Then, an Ion Xpress Plus Fragment Library Kit (Life Technologies) was employed to prepare barcoded shotgun libraries. To conduct emulsion PCR, an Ion PGM Template OT2 400 Kit (Life Technologies) was utilized. Sequencing was performed by the Ion Torrent PGM (Life Technologies) analyser using Ion 318 chips and Ion PGM sequencing 400 Kit v2 (Life Technologies) in accordance with the manufacturer’s protocol.

### Transcriptome assembly and annotation

Cutadapt [95] <u>v1.9</u> was applied to trim the adapter sequences used for cDNA synthesis. Prinseq lite [96] v.0.20.4 was used to trim reads according to their quality and length. For *de novo* transcriptome assembly, we used Trinity software [20] (version r20131110) with the default parameters. Blast2GO [21] was used for Gene Ontology (GO) analysis and the functional annotation of contigs. Local BLAST (BlastX threshold value of e = 1×10^-6^, matrix BLOSUM-62) and the nr database were used for the grouping and annotation of contigs. MEGAN6 software [93] was used to visualize the contig distribution (by KOG/EGGNOG classifications). TransDecoder and Trinotate were applied to identify and annotate ORFs.

### Differential expression analysis

Differentially expressed genes were detected according to a recent protocol using the Bowtie [91], Htseq [97] and edgeR [98] software packages. Because biological replicates were available only for *H. verbana,* we applied the dispersion estimates for *H. verbana* to other leech species. Genes with q-value (FDR) < 0.05 were defined as differentially expressed. To perform differential expression analysis using a genome model, the cDNA reads of *H. medicinalis* were mapped against the genome assembly using the STAR software [99]. In addition to the cDNA reads corresponding to the salivary cells and muscles, we also mapped reads of *H. medicinalis* ganglion 2 against its genome assembly (SRR799260, SRR799263, SRR799266). The HTSeq software [97] was used to count the number of reads mapped against the annotated genes. For each tissue (salivary cells, muscles, and neural tissue), we established a list in which the numbers of mapped reads corresponded to individual gene IDs by applying the “htseq-count” python script with the default parameters. Unique transcripts corresponding to certain tissues were determined by finding the intersection of these lists.

### Collection and concentration of the salivary cell secretions

To collect SCSs from three medicinal leech species, the bottom of a 15 mL Falcon tube was cut off, and an impermeable membrane (PARAFILM^®^ M) was stretched on the excised end. The tube was filled with saline solution containing 10 mM arginine. A leech bit through the membrane, sucked up the salt solution and emitted its secretion into the saline solution. The saline solution enriched with SCS was continuously stirred and renewed to prevent its ingestion by leech. We collected approximately 10 mL of saline solution containing highly diluted leech SCS. Harvested SCSs were concentrated on a solid-phase extraction Sep-Pak Vac C18 6cc cartridge (Waters, USA) using 0.1% TFA in 70%/30% acetonitrile/water (v/v) as the buffer for elution of the protein fraction. The acetonitrile was evaporated, and the remaining solution was lyophilized. Dry protein powder was stored at −70°C prior to mass spectrometric analysis.

### Digestion of the salivary cell secretions and sample preparation for proteomic analysis

Several digestion and preparation methods were applied to each freeze-dried sample of the SCS to cover the broadest possible variety of the salivary proteins. These preparation methods included filter-aided sample preparation (FASP), gel-free trypsin digestion using surfactant RapiGest SF (Waters), and ingel digestion of the protein sample. Detailed descriptions of the preparation methods are presented in the **Supplementary Methods**.

### Mass spectrometry and analysis of mass spectra

Mass spectra were obtained for each prepared protein sample by using three instruments: (i) a TripleTOF 5600+ mass spectrometer with a NanoSpray III ion source (Sciex, USA) coupled to a NanoLC Ultra 2D+ nano-HPLC system (Eksigent, USA), (ii) a Q-Exactive HF mass spectrometer with a nanospray Flex ion source (Thermo Scientific, Germany), and (iii) a Maxis 3G mass spectrometer with an HDC-cell upgrade and an Online NanoElectrospray ion source (Bruker Daltonics GmbH, Germany) coupled to a Dionex Ultimate 3000 (ThermoScientific, USA) HPLC system. The obtained raw mass spectra were converted into non-calibrated peaklists by the appropriate software, and these peaklists were analysed using ProteinPilot 4.5 revision 1656 (ABSciex). The acquisition of the mass spectra and their analysis and peptide identification are described in detail in the **Supplementary Methods**.

To create a final list of the identified proteins for each medicinal leech species, the combined transcriptomes were translated either by ORF Transdecoder (standard parameters) or by the six-reading-frame method (length filter was ≥30 aa). Then, the protein sequences obtained by both methods were combined and used to establish a referential database of potential SCS proteins for each medicinal leech species. The protein sequences in the referential database that were matched by the peptides identified in an individual peaklist allowed the creation of protein datasets for individual samples. The protein datasets generated for individual samples by using different preparation methods and mass spectrometry techniques were combined to create the final list of the SCS proteins for each medicinal leech species.

## Data Access

All of the raw reads generated in this study have been deposited in the NCBI database under BioProject accessions PRJNA257563 and PRJNA256119.

## Acknowledgments

*H. medicinalis* genome sequence, assembly and annotation were supported by the Russian Science Foundation (project No17-75-20099).

Transcriptomic analysis of salivary and muscle cells of *H. medicinalis, H. verbana, H. orientalis* and proteomic analysis of their salivary cell secretions were supported by the Russian Science Foundation (project No14-14-00696).

O.V. Podgorny was partially supported by the IDB RAS Government basic research program, no. 0108-2018-0007. Laser microdissection was performed using equipment of the Core Centrum of Institute of Developmental Biology RAS.

V.G.Z mass spectrometric measurements were performed using the equipment of “Human Proteome” Core Facility of the Orekhovich Institute of Biomedical Chemistry (Russia) which is supported by Ministry of Education and Science of the Russian Federation (agreement 14.621.21.0017, unique project ID RFMEFI62117X0017)

M.S.G. is grateful to the Russian Science Foundation (grant No. 14-50-00150) for support.

We thank, Alexander V. Bazin the director of the company HIRUD I.N. Ltd.

## Author contributions

V.N.L., O.V. Podgorny and V.V.B. conceived and designed the experiments. I.P.B., V.A.M., A.S. Kurdyumov, and D.A.S. samples collection. O.V. Podgorny, D.A.S. and V.V.B. laser microdissection.

V.V.B., D.A.S., and V.N.L. generated DNA and RNA for sequencing. V.V.B., M.T.V. and E.S.K. genome and transcriptome sequencing. V.V.B., A.S. Kasianov, and A.I.M. genome assembly, binning and automated annotation. V.V.B., A.S. Kasianov, and D.V.V. transcriptome assembly and annotation. D.V.V., A.S.N. differential expression analysis. V.G.Z., S.I.K., N.A.A., I. O.B., O.V. Pobeguts, D.S.M and D.V.R. sample preparation and mass spectrometry for proteomic analysis. V.V.B., E.N.G. analyzed results and made the figures. V.V.B. and O.V. Podgorny wrote the paper. I.P.B., M.S.G., V.M.G., H.B.S. and V.N.L. coordinated and revised the paper.

